# The plant specific VPS2.2/HYADE of the ESCRT-III protein sorting machinery localizes to a secretory compartment and is required for cell division plane determination in Arabidopsis

**DOI:** 10.64898/2026.07.15.738706

**Authors:** Verena Ibl, Christian Pritz, Verena Köhler, Sabine Müller, Nikolaus Gössweiner, Ilse Foissner, Marie-Theres Hauser

**Affiliations:** University of Natural Resources and Life Sciences, Muthgasse 19, 1190 Vienna, Austria; University of Applied Sciences Dresden (HTW Dresden), Pillnitzer Platz 2, 01326, Dresden, Germany; Universitä t Salzburg, Hellbrunnerstr. 34, 5020 Salzburg; NICO – Neuroscience Institute Cavalieri Ottolenghi Azienda Ospedaliero – Universitaria San Luigi Gonzaga Regione Gonzole, 10 – 10043 Orbassano; Friedrich-Alexander-Universitä t, Department of Biology, Staudtstr. 5, 91058, Erlangen, Germany; Europagymnasium Auhof, Aubrunnerweg 4 4040 Linz, Austria; Universität Salzburg, Hellbrunnerstr. 34, 5020 Salzburg

**Keywords:** cytokinesis, secretory pathway, ESCRT, Arabidopsis, roots, HYADE

## Abstract

Positioning of the division plane is a critical step during cellular development. While in animal cells the plane of cell division is defined during metaphase, in plants this decision is made before mitosis and includes the formation of a cytoskeletal structure termed the preprophase band (PPB). Although not essential, the PPB constitutes the earliest mark for the plane of cell division and coincides with the sites where the cell plate fuses with the parental plasma membrane during cytokinesis. Recent studies indicate that endo- and exocytosis are involved in remodelling of the cortical division zone. Here we show that PPBs, phragmoplasts, cell plates and cell walls are misplaced in Arabidopsis mutants of *Atvps2.2/hyade (hya)*, a plant-specific component of the Endosomal Sorting Complex Required for Transport (ESCRT)-III. We demonstrate that the amino acid substitution Q71P of the *hya-3* allele abolishes the homo- and heteromerization with other ESCRT-III components. Functional HYA-GFP fusion proteins are excluded from the canonical location of ESCRT-III complexes at multivesicular bodies (MVBs) and are absent from Brefeldin A (BFA) and Wortmannin (WM) sensitive compartments in *Arabidopsis*. HYA-GFP neither localize to early nor late endosomes but colocalizes with markers of secretory compartments such as the Qc-SNARE, SYP61 and with trans-Golgi-network (TGN) derived secretory Rab-A3 vesicles. Moreover, HYA-GFP locates to the extracellular space indicating that VPS2.2/HYA is involved in secretion. These results are consistent with a novel, non-canonical function of ESCRT-III in the establishment and maintenance of the cortical division zone via secretion.

## INTRODUCTION

During plant cell division, the spatial, structural, and molecular organization of the cortical division zone (CDZ) controls the division plane determination^1–4^. Prior to entry into prophase, the PPB, a transient cortical microtubule and actin structure, is formed at the cortical division zone and determines the plane of cell division^5^. The PPB leaves behind an unknown mark that guides the expanding cell plate toward the pre-determined division zone during cytokinesis. Upon break-down of the PPB a region complementary to the CDZ is formed by apico-basal actin enrichment or pronounced reduction of actin, often times referred to as actin depleted zone (ADZ) and reduction of kinesin-like protein KCA1 depletion zone (KDZ)^2,6^. Furthermore, several proteins play a role in division plane determination and plasma membrane modification, which are deposited at the cortical division zone via vesicle trafficking: TPLATE, that functions in vesicle-trafficking events required for site-specific cell wall modifications; the microtubule structure binding protein AIR9; the HRGP-type cell wall protein RSH, which functions in the correct positioning of the cell plate^5^; TANGLED1, that binds to PPBs, spindle and phragmoplasts and is involved in cell plate guidance together with AIR9^7^. Furthermore, RanGAP1, which is recruited to the PPB and cortical division site in a dephosphorylated (FASS/TONNEAU 2)- and kinesin (POK1/POK2) dependent manner has a role in the spatial signalling during plant cell division^3^. Even though early cell division mechanisms differ between plant and animal cells, the late stages of cytokinesis appear to be similar^8,9^. The phragmoplast, which emanates from the spindle midzone and is essential for the formation of the cell plate, resembles the midbody, sharing its structural and molecular features^10^. During the cell plate assembly, Golgi- and TGN-derived vesicles, and vesicle-associated molecular machineries like Rab GTPases (Rab-A2, Rab-A3, and RAB-A1d), SNARE proteins, dynamin-related proteins, EXOCYST tethering complexes and actin are responsible for the delivery, tethering and fusion of vesicles at the cell plate assembly matrix (CPAM) and subsequently for the cell plate biogenesis^11^. Furthermore, fusion of cell plate and plasma membrane and their heterotypical fusion are crucial for the completion of cytokinesis. This membrane fission and fusion is topologically similar to budding events. Studies of the ESCRT machinery in yeast and animal cells have revealed that the assembly of ESCRT-III polymers is important for unrelated biological processes by mediating multivesicular bodies (MVBs) biogenesis, cytokinesis and retroviral budding by membrane budding away from the cytosol^12,13^. Moreover, a few examples for a key role of the ESCRT system in cytokinesis have been provided: the ESCRT-I component Tsg101/VPS23 and an ESCRT-associated protein ALIX are recruited to the midbody^14–16^, the ESCRT-III component CHMP1B binds and recruits the microtubule-severing enzyme spastin to the midbody^17^; and ESCRT-III components and VPS4 in Archaea are localized to the mid-cell of dividing cells^18,19^. In Arabidopsis, VPS23/Tsg101 ELCH, is described as a putative key regulator of the microtubule cytoskeleton during cytokinesis^20^. Dominant negative AtVPS4/SKD1 mutant lines were described with an increased amount of trichomes with multiple nuclei and more branches^21^. Here, we show that HYA is part of a secretion mechanism during plant cell division that is necessary for plasma membrane remodelling.

## RESULTS

### *hya* is defective in cytokinesis in Arabidopis root

*Hyade* (*hya*) was isolated as a cytokinesis defective multinucleated Arabidopsis mutant with incomplete cell walls^1^ (Fig. 1a-h). Aberrantly positioned transverse cell walls lead to misshaped cells in all four *hya* alleles (Fig. 1f, g). Previously performed immunolocalization studies showed that PPB and phragmoplasts are tilted^1^. To quantify *in vivo* the involvement of HYA in positioning of these microtubule arrays, the microtubule marker line GFP-MAP4^22^ was introduced to *hya-1* and *hya-3* (Fig. 1i-k). The deviation from the midline was used as measure for the extent of misalignment and is pronounced in PPBs and even more severe in phragmoplasts (Fig. 1l), indicating that in *hya* mutants the cell-plate navigation and tilt-correction^23^ is reduced. Since secretory and endocytic trafficking events are proposed^23,24^ to be involved in the establishment of the division plane and the formation of the cell plate in plants, endocytosis was quantified by pulse labelling of root epidermal cells with the *in vivo* dye FM4-64^25,26^. The plasma membrane of *hya-4* was loaded significantly less with FM4-64 compared to wild-type which may point to an altered membrane composition. Additionally, the FM4-64 fluorescence disappeared faster in *hya-4* (Fig. 1m-o) than in wild-type. Although, in *hya-4* as well as in wild-type, FM4-64 finally reaches the vacuolar membrane (data not shown). This demonstrates the functional endocytic trafficking. Since plant ESCRT-III proteins are described to function on the MVB to insulate membrane cargos and to generate intraluminal vesicles (ILV), we analyzed if the morphology of MVBs is affected in the mutant line. We found that the MVB diameter, and the number of ILVs in *hya-3* are not altered compared to wild-type ones (Suppl. Fig. 1a, b). Concerning the ILV diameter, we noticed a slightly smaller diameter of MVBs in *hya-3* (Suppl. Fig. 1a, b). Since the scatter plot indicates only a small shift to fewer numbers of MVBs with a smaller diameter in *hya-3* compared to wild-type (Suppl. Fig. 1a, b), we conclude that the biogenesis of MVBs is not the main functional target of HYA.

**Figure 1.**
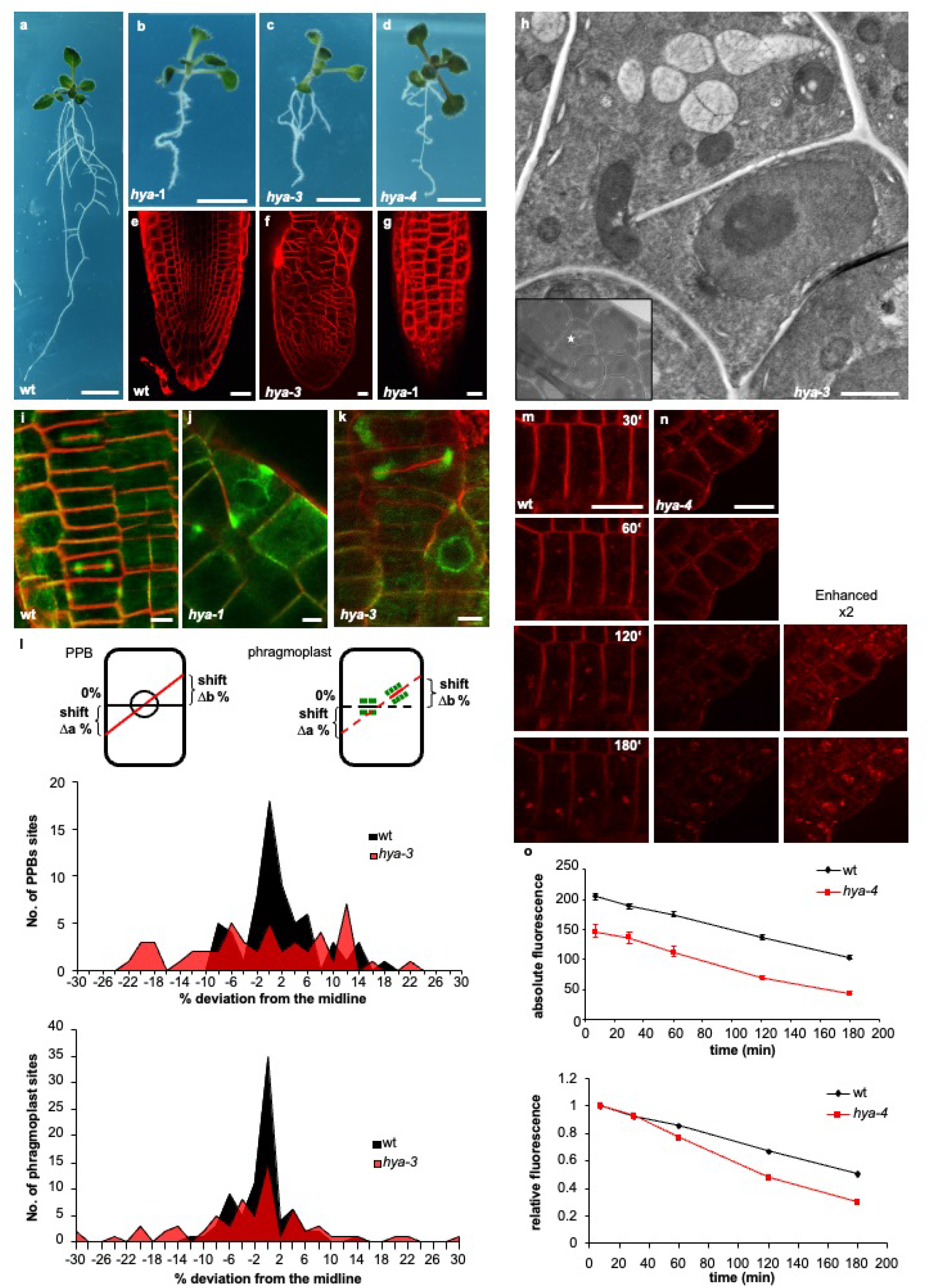
Root specific phenotypes of *hya* mutants. 11 day old wild-type (a), *hya-1* (b), *hya-3* (c) and 13 day old *hya-4* (d) seedlings. The meristem architecture was visualized with FM4-64 of wild-type (wt) (e), *hya-3* (f) and *hya-1* (g). (h) Electron micrographs of a *hya-3* cell with an incomplete cell wall. Insert displays an overview of magnified area in the meristem. PPBs (i, j, k) and phragmoplasts (i, k) in wt (i), *hya-1* (j) and *hya-3* (k), respectively. (l) Quantification of the extent of misaligned PPBs and phragmoplasts. The graphs summarize the deviation from the midline of 32/25 wt/mutant PPBs and 41/35 wt/mutant phragmoplasts. Endocytosis was quantified at the plasma membrane with a time course of FM4-64 internalization on wt (m) and *hya-4* (n) epidermal cells of 7-8 day old seedlings. (o) The graphs show the absolute and relative fluorescence at plasma membranes of 260 wt (black) and 141 *hya-4* (red) cells. The relative fluorescence was normalized to the signal directly after FM4-64 incubation and illustrates the difference between the velocity of endocytosis in the first 30 min and at later time points. Bars indicate ± SE (standard error) from averages of five independent experiments. Scale bars represent 10 mm (a-d), 20 µm (e-g), 1 µm (h), 6 µm (i-k) 15 µm (m) and 30 µm (n).

### HYA/VPS2.2 clusters to a plant specific ESCRT-III brunch

The (*HYA*) gene was cloned by a map-based approach (Suppl. Fig. 2a-c) and is synonymous with the ESCRT-III component AtVPS2.2 (At5g44560)^1^. *hya-1* and *hya-2* have identical deletions in exon 7, *hya-3* has a missense mutation in exon 5 that leads to a substitution of the polar amino acid Glutamine (Q) to Proline (P) at residue 71 and in *hya-4* the T-DNA is inserted in exon 6 (Suppl. Fig. 2b). The identity of the (*HYA*) gene was confirmed by complementation of *hya-2* with a genomic 3.3 kb fragment of the At5g44560 gene (Suppl. Fig. 2c). We used protein queries to search translated nucleotide databases (TBlastN analyses) to evaluate the presence of the plant specific branch of VPS2 in the ESCRT-III subcomplex^27^. We included mammals, fishes, insects, green algae, land plants, and ascomycetes. HYA (AtVPS2.2) clusters in the land plant specific branch VPS2.2 in cluster VPS2A (Fig. 2a). Interestingly, no plants and no ascomycetes were found for VPS2B, suggesting specificity and confirm that complex gene duplication is existing for ESCRT-III complexes^28^. Moreover, these findings suggest that HYA is possibly involved in a distinct mechanism for ESCRT-III complexes. Although HYA clusters in a plant specific lineage of VPS2, we described recently that AtVPS2.2 interacts with the core (AtSNF7, AtVPS20) and coat (AtVPS2, AtVPS24) ESCRT-III components and with associated proteins such as AtVPS4, AtVPS60.1 and AtVPS46.1/2^29^ which is summarized as schema in Fig. 2b. All ESCRT-III components are similar in size, share six predicated alpha-helices, two in the basic N- and four in the acidic C-terminal half^30^ and have a crystal structure similar to the human HsVPS24 (CHMP3)^31^. To map the effect of the *hya-3* mutation its 3-D structure was modelled to the CHMP3^31^ using Chimera^32^. The protein alignment revealed that the Q71P substitution is located in the middle of helix *α*2 (Fig. 2c) which forms the 70 long hairpin critical for membrane binding and homo- and heterodimerization of ESCRT-III components. Additionally, the residues in *α*2 and *α*4 face the nonpolar bilayer^13^ (Fig. 2c). Using Chimera, we predicted the Q71P mutation which produced clashes with L68 and A67 within *α*2, putatively inducing a deformation of helix *α*2. Additionally, we used AlphaFold^33,34^ to run homo- and heterodimerization prediction of HYA/HYA, *hya-3*/HYA, *hya-3*/*hya-3*, HYA/AtVPS46.1/2 and *hya-3*/AtVPS46.1. Whereas the predicted HYA/HYA homodimer is parallel (Fig. 2d), the homodimer of *hya-3*/HYA is antiparallel (Fig. 2d) and the homodimer of *hya-3*/*hya-3* the parallel arrangement is affected (Fig. 2d). In addition, *hya-3*/AtVPS46.1/2 showed an altered 3-D structure compared to HYA/AtVPS46.1/2 (Fig. Suppl. 3). Yeast two-hybrid (Y2H) analysis of *hya-3* with HYA, AtVPS46.1/2 and *hya-3* proofed that the substitution altered the interaction strength (Fig. 2e). These data suggest, that the Q71P mutation completely abolishes the function of HYA/VPS2.2 and may explain the similarity of the *hya-3* phenotype with knock out alleles.

**Figure 2.**
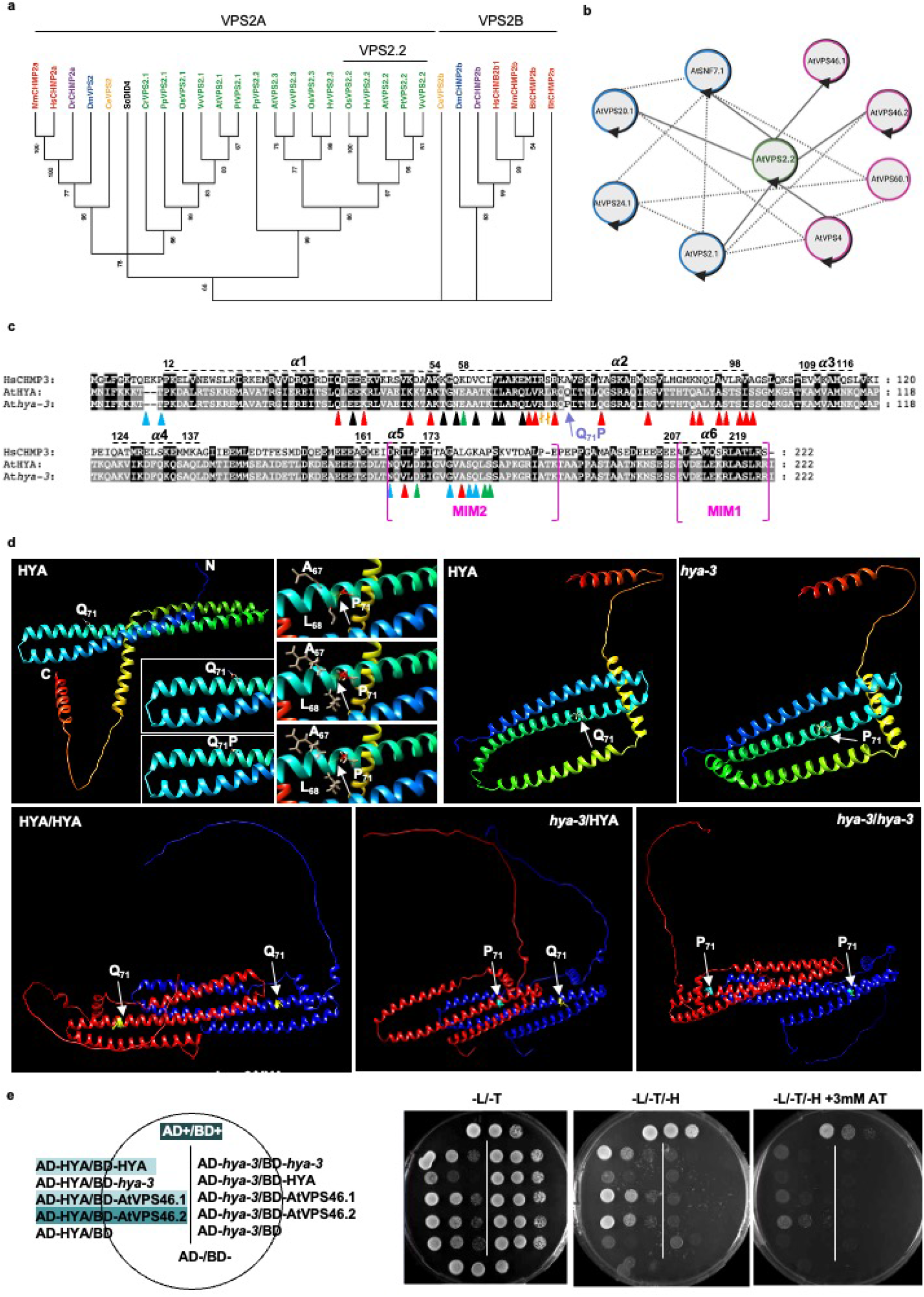
Characterization of VPS2.2/HYA. (a) Phylogenetic reconstruction of VPS2 ESCRT family proteins constructed by the software Molecular Evolutionary Genetics Analysis (MEGA) X^62^. Bootstrap was set to 1000. The first two letters of each protein name represent the organism and the nomenclature is colour coded like following: red: Mm, *Mus musculus*; Hs, *Homo sapiens*; violet: Dr, *Danio rerio*; blue: Dm, *Drosophila melanogaster* ; orange: Ce, *Caenorhabditis elegans*; black: Sc, *Saccharomyces cerevisiae*; green: At, *Arabidopsis thaliana*; Pt, *Populus trichocarpa*, Vv, *Vitis Vinfera*; Os, *Oryza sativa*; Pp, *Physcomitrella patens*; Cr, *Chlamydomonas reinhardtii*, Hv, *Hordeum vulgare*. (b) Schematic representation of the known interactions between AtVPS2.2 (green), ESCRT-III proteins (blue) and ESCRT-III associated proteins (pink) indicated in grey lines. All proteins show homo-dimerization (arrow) except AtVPS60.1. Interactions between all these proteins in indicated with dashed lines. (c) Structure based-protein-sequence alignment of human HsCHMP3, HYA and *hya-3*. The Q71P substitution in *hya-3* is indicated in purple and is in helix *α*2. In HsCHMP3 the same position is occupied by a helix stabilizing alanine (A) whereas P is a known helix disruptor. Important residues are indicated according to^31^: blue triangles: residues involved in dimer-dimer interactions (concave dimer); red triangles: residues involved in CHMP3 dimerization; black triangles: residues involved in dimer-dimer interactions (dimer 2); and green triangles: residues involved in dimer-dimer interactions (convex dimer). The magenta brackets indicate the ESCRT-III conserved MIM1 (Obita, Saksena et al. 2007) and the MIM2 (Kieffer, Skalicky et al. 2008) domain. (d) The structure prediction of *hya-3* shows that the Q71P substitution locates in helix *α*2. Structure editing analysis performed with Chimera (Pettersen, Goddard et al. 2004) identifies clashes with L68 and R67, an area, that is important for dimerization (Muziol, Pineda-Molina et al. 2006). Alpha-Fold prediction of HYA/HYA, *hya-3*/HYA and *hya-3*/*hya-3* homodimerization shows an altered structural pattern. (e) Growth assays from HYAHYA, HYA/hya-3, HYA/AtVPS46.1/2 are shown. The identity of the co-transformants is indicated on the left, the selective media above and the strength of the interactions on the right. Shown are spotted undiluted and 1:10 diluted cultures of co-transformants. -l/-t: media without leucine, tryptophan; -l/-t/-h: media without leucine, tryptophan, histidine; 3-AT, 3-Aminotriazole

### HYA is strongly expressed in roots and localizes to mobile intracellular compartments of different sizes

To follow HYA expression in different tissues, we used the *β*-glucuronidase (GUS) reporter (pmHYA::GUS). Two days after germination GUS staining was first detectable in the root meristem and vasculature (Suppl. Fig. 4a-d). In the vasculature of rosette leaves and flowers, pmHYA::GUS expression was only detectable after prolonged staining in the stigma, style and filaments of flowers (Suppl. Fig. 4e,f). Subcellular localization of HYA was visualized in wild-type, and in *hya-1* - and *hya-3* plants transformed with the functional construct pmHYA::gHYA-GFP (gHYA-GFP) as previously described^29^ (Suppl. Fig. 4h, i). *hya-3* was complemented by gHYA-GFP and its expression was strong in the meristem and elongation zone of roots (Suppl. Fig. 4h, i). Notably, the expression of pUBQ10::YFP-gHYA did not rescue *hya-3*, supporting that gHYA-GFP is functional and indicating an affected N-terminus of the pUBQ10::YFP-gHYA construct (Suppl. Fig. 4i,j). Plant ESCRT proteins have been described to be involved in the formation of MVBs^35^. Since *hya-3* does not affect the morphology of MVBs (Suppl.Fig. 1), we speculate, that HYA is localized at an additional endosomal compartment to MVBs. We monitored the expression in the root meristem and observed gHYA-GFP at globular intracellular mobile compartments of different sizes, as well as in predominantly transverse oriented plasma membranes (Fig. 3a). To test whether the inhibition of vesicle trafficking and the blocking of endocytosis and MVB biogenesis is affecting the localization pattern of gHYA-GFP, we treated seedlings expressing gHYA-GFP with Brefeldin A (BFA)^36,37^ and Wortmannin (WM)^24,38^, respectively. Previously, it was shown that Arabidopsis SORTING NEXIN 1 (AtSNX1) localizes to an endosomal compartment sensitiv to both, BFA and WM^39^. While AtSNX1-GFP vesicles assembled to larger aggregates already after 30 min of BFA exposure (Fig. 3b), the motility, shape, and size of gHYA-GFP compartments were unaffected (Fig. 3c). Similarly, WM treatment led to enlarged, vacuolated AtSNX1-GFP compartments but did not alter gHYA-GFP localization (Fig. 3d, e). Next, we analyzed the cytoskeletal requirements for the guidance of gHYA-GFP-containing vesicles. We interfered with the cytoskeleton formation using the actin and microtubule-depolymerizing agents latrunculin B (Lat B)^40^ and oryzalin^41^, respectively. Lat B treatments disrupted F-actin in the GFP-FABD2^42^ transgenic actin reporter line (Fig. 3f), reduced the mobility of (larger) gHYA-GFP agglomerated compartments (Fig. 3g, Supplementary Information, Movie 1) and stopped mobile FM4-64 labeled compartments (Fig. 3h, Supplementary Information, Movie 2, 3). Solely, the small gHYA-GFP vesicles remained unaffected (Fig. 3g, Supplementary Information, Movie 1). By contrast, oryzalin treatment clearly affected microtubules of the GFP-MAP4 transgenic line (Fig. 3i) but did not visibly interfere with gHYA-GFP trafficking (Fig. 3j, Supplementary Information, Movie 4) and FM4-64 labeled endosomes (Fig. 3k, Supplementary Information, Movie 5, 6). These results confirm that plant endosomal trafficking is mostly actin–dependent^36,43,44^, and revealed that gHYA-GFP positive compartments are hardly affected by cytoskeleton inhibitor treatment. Furthermore, colocalization of gHYA-GFP with FM4-64 vesicles was never observed during these experiments (Fig. 3a-k). Following the signal of gHYA-GFP during cytokinesis revealed that small gHYA-GFP-containing vesicles are moving to the emerging cell plate (Fig. 3l, Supplementary Information, Movie 2). Larger agglomeration, which are formed by smaller ones (data not shown) are moving to the cortical division site at the late stage of cytokinesis (Fig. 3l). To verify these localizations, we studied gHYA-GFP localization in transiently transformed protoplasts. Notably, gHYA-GFP localized around the FM4-64-stained cell plate and further accumulated at the cortical division site during cytokinesis (Suppl. Fig. 5). Since hya shows already a pre-mitotic effect on the PPB, we presumed that HYA is located at the cortical division site during preprophase. It is worth to mention, that we monitored within seven cytokinesis events four preprophase events, where HYA localizes at the cortical division site.

**Figure 3.**
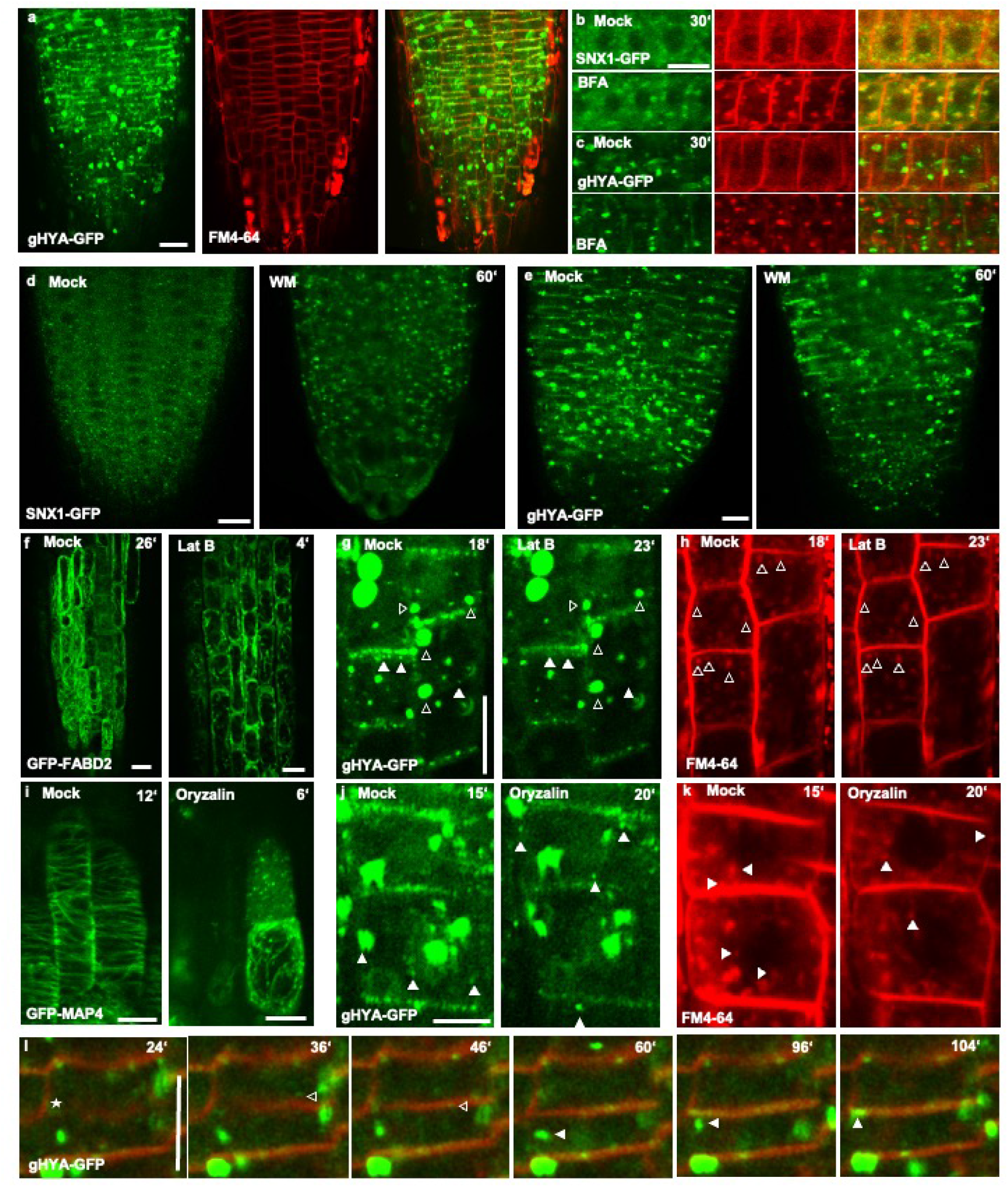
HYA is BFA- and WM-insensitive and localizes to novel vesicular compartments and at the cell plate. (a) gHYA-GFP defines compartments of different size and localizes predominantly in transverse plasma membranes. (b, c) SNX1-GFP (b) but not gHYA-GFP (c) shows BFA sensitivity after 30 min of 100 µM BFA treatment. (d, e) SNX1-GFP and gHYA-GFP WM-treated with 100 µM for 1 h. Note that gHYA-GFP is not affected by WM treatment (e). (f-h) Inhibition of polymerization of actin by Lat B (10 µM) shows F-actin depolymerization defects after 4 min treatment in root tips expressing FABD-GFP (f). In contrast to small gHYA-GFP positive vesicles (filled arrowheads), agglomerated gHYA-GFP positive vesicles (open arrowheads) - as well as FM4-64 are sensitive to Lat B (g, h). (i-k) Depolymerization of microtubules with oryzalin (10 µM) in a GFP-MAP4 transgenic line (i). By contrast, there was no effect on gHYA-GFP and FM4-64 vesicles (filled arrowheads), respectively (f-k). Time of starting- and endpoint of the 5 minutes movie is indicated. (l) gHYA-GFP vesicle trafficking in stable transformed Arabidopsis root cells during cytokinesis. Small gHYA-GFP positive vesicles (open arrowheads) are moving to and localizing around the cell plate (asterisk) at later stages of cytokinesis. Larger vesicles (filled arrowheads) are moving to the cortical division site after fusion of the cell plate with the plasma membrane. Time section represents 120 min of a movie, specific timepoint is indicated. Scale bars represent 10 µm (a), 15 µm (b-c), 10 µm (d-e), 30 µm (f-h), 20 µm (i-k, l).

### HYA to TGN-derived compartments and vesicles involved in exocytosis, but not to MVBs

Our inhibitor treatments indicate that gHYA-GFP positive compartments are distinct from known endosomal compartments. To further confirm this notion, we performed first immunolabeling experiments of high-pressure frozen gHYA-GFP Arabidopsis roots using anti GFP antibody. gHYA-GFP is localized at the plasma membrane and at vesicles resembling prevacuolar compartments (PVC) (Fig. 4a). We continued with double immunolabeling experiments using the characterized antibodies against AtVSR1^45^ that label both, MVBs and PVC. Under our experimental conditions, gHYA-GFP colocalized with the native AtVSR1 protein at PVCs (Fig. 4b-d). In contrast to the AtVSR1 localization, we observed gHYA-GFP near the plasma membrane (Fig. 4a) but never in MVBs (n = 75; Fig. 4d). It is known that transgenic GFP-SKD1/VPS4 Arabidopsis seedlings showed colocalization of FM4-64 and GFP-SKD1 positive compartments, suggesting that GFP-SKD1 is targeted to an endosomal multivesicular compartment^46^. However, immuno gold labelling and electron microscopy studies confirmed the inhibitor treatments, suggesting that HYA associates with compartments that are distinct from known endosomal compartments. Next, we analysed if HYA is co-localizing with distinct proteins involved in endocytic and secretory trafficking. For this, we generated the construct 35S::mCherry-HYA that co-localises with gHYA-GFP in transiently transformed mesophyll protoplasts (Suppl. Fig. 5a-c). Subsequently, we co-transformed 35S::mCherry-HYA together with soluble N-ethylmaleimide-sensitive factor (NSF) attachment protein receptors (SNAREs) and RAB reporter proteins. We did not observe co-localization of 35S::mCherry-HYA with the late endosomal marker ARA6-GFP^47^ in mesophyll protoplasts, confirming the immunolabeling result, that HYA does not localize to MVBs (Fig. Suppl 5d-f). Additionally, no colocalization was identified of HYA with the early endosomal marker GFP-ARA7^47^ (Fig. Suppl. 5g-i). However, HYA showed partial co-localization with the cell-cycle dependent and plant-specific Qc-SNARE protein SYP71^48^ which is supposed to localize at the endoplasmic reticulum (ER), at the plasma membrane (PM) and endosomes^49^ (Fig. 4e-j). Additionally, HYA showed co-localization with RAB-A3, at the TGN-derived RAB-A2/A3 compartment which is part of the secretory pathway^50^. These results indicate, HYA associates with endomembrane compartments of the secretory system. To assess the localization of HYA on the secretory system, we analysed the localization of HYA during cell division in mesophyll protoplasts. Indeed, gHYA-GFP appears to localize outside of the plasma membrane at later stages of cytokinesis, indicating possible localization at the cell wall (Fig. 5a). To confirm the localization of HYA at the cell wall, we induced plasmolysis of gHYA-GFP expressing root cells. gHYA-GFP localizes around the plasma membrane as well as at the cell wall (Fig. 5b, c). No signal was found at Hechtian strands (Fig. 5b), indicating that HYA is part of some secretion mechanism.

**Figure 4.**
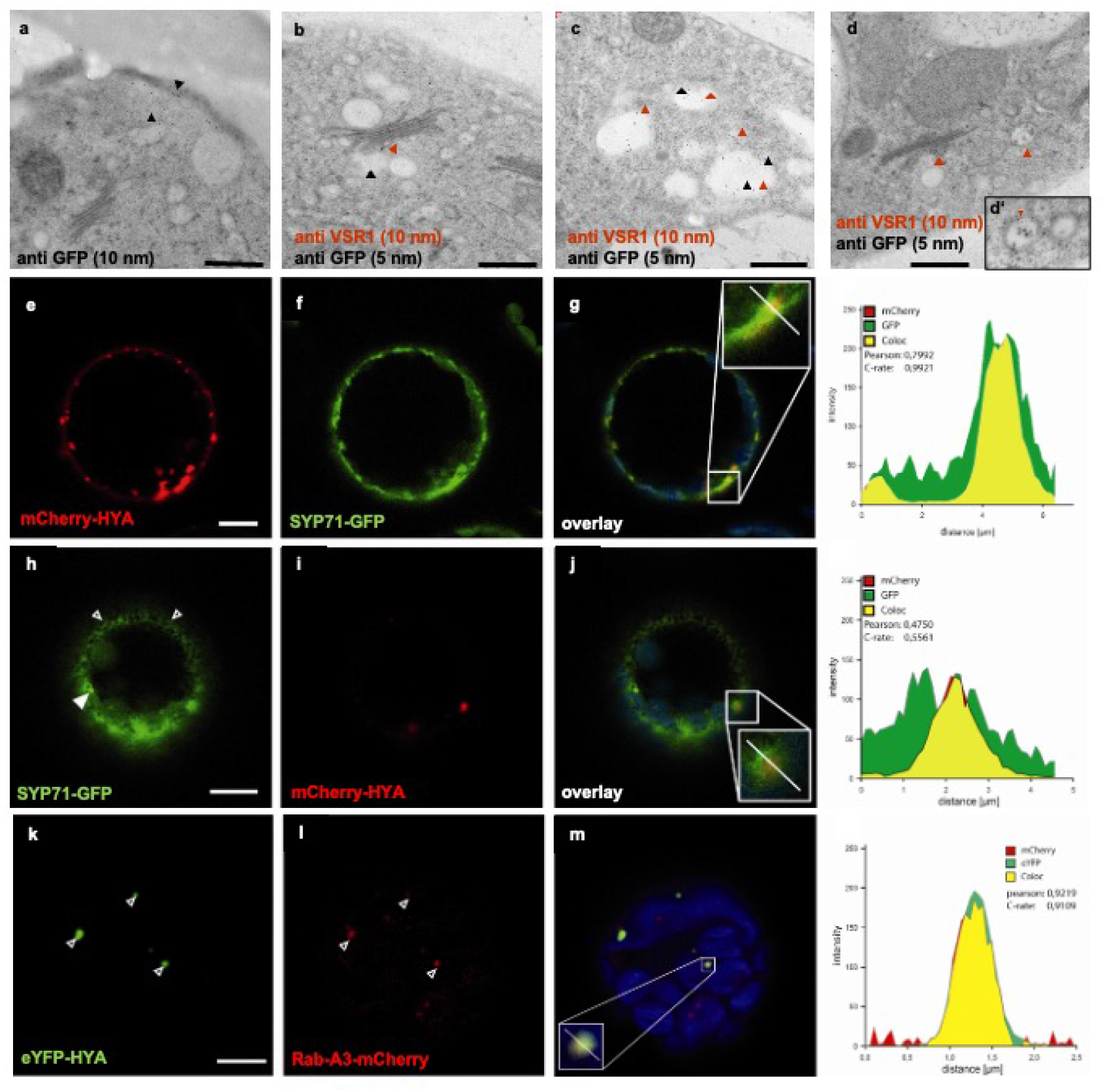
HYA localizes to PVC compartments and to secretory vesicles. (a-d) Immunolabelling with anti-GFP alone (a) and together with anti-VSRAt-1 (b-d). Black arrows indicate anti-GFP gold particles at the plasma membrane and in vesicles resembling PVCs. Anti-VSRAt-1 (red arrows) labels TGN (b, d) and MVBs (n = 4; d, d’). (c) Partial colocalization of gHYA-GFP was detected in PVCs. (e-j) Partial co-localization of mCherry-HYA with SYP71-GFP. (k-m) Co-localization of eYFP-HYA with RabA3-mCherry. Note the fluorescence profiles, respectively. Scale bars represent 0.5 µm (a-d’), and 10 µm (e-m).

**Figure 5.**
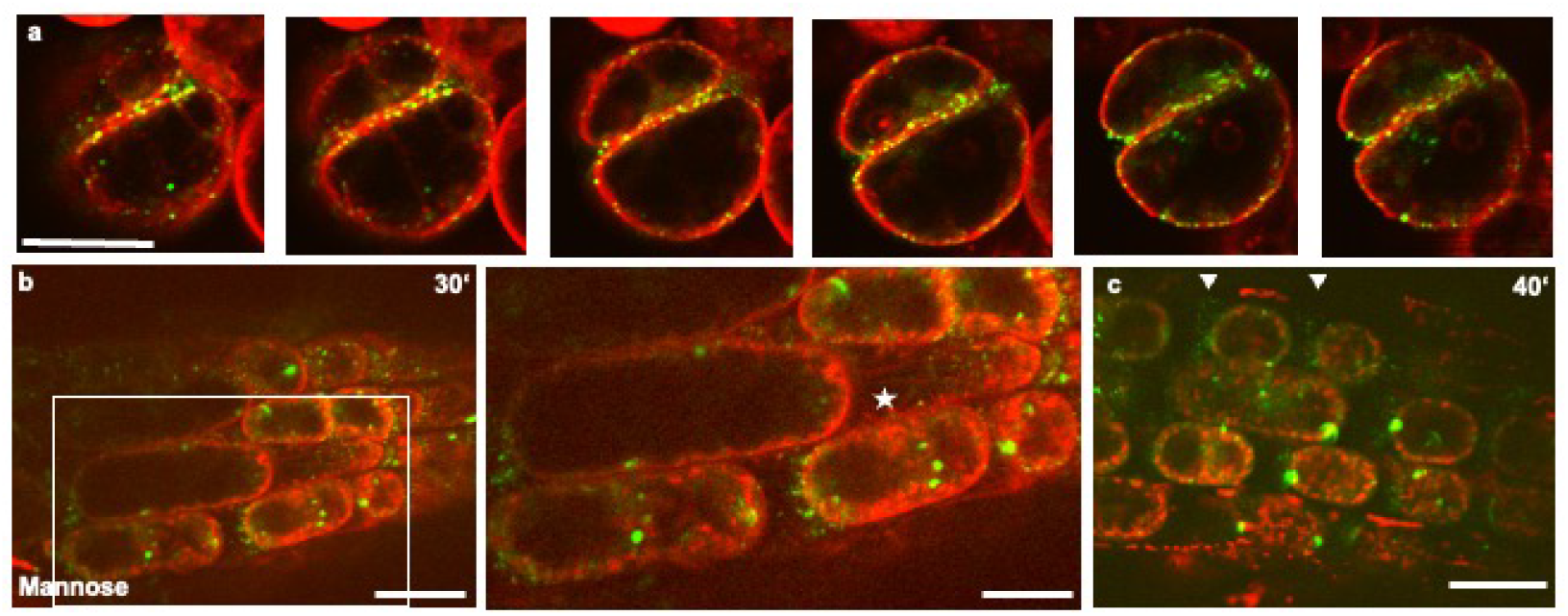
gHYA-GFP localization during late cytokinesis and after plasmolysis treatment. (a) gHYA-GFP transformed protoplasts were incubated over night at room temperature in darkness and stained in a 16 µM FM4-64 solution for 3 to 5 hours. After completing cell division gHYA-GFP localizes outside the cell membrane. (c) gHYA-GFP vesicle trafficking in stable transformed Arabidopsis root cells during cytokinesis. Small gHYA-GFP positive vesicles (filled arrowheads) are moving to and localizing around the cell plate at later stages of cytokinesis. Larger vesicles are moving to the cortical division site after fusion of the cell plate with the plasma membrane. Time section represents 120 min of a movie, specific timepoint is indicated. (b, c) gHYA-GFP localization in 7 day old root cells after 30 (b) and 40 min (c) of mannose treatment. After 20 min incubation, the tonoplast was separated from the cell wall. Asterisks mark the hechtian strands, where no gHYA-GFP signal was observed. Arrows indicate HYA localization on the cell wall. Plasma membrane was stained 4 min with 8 µM FM4-64, plants were mounted in 0.5 M mannose solution. To visualize Hechtian strands, the contrasts of the pictures were enhanced. Scale bars equal 20 µm (a) and 30 µm (b, c).

### Conclusions

Here, we have identified HYA as an indispensable plant specific ESCRT-III component that is necessary for the correct positioning of the cell division plane in the root meristem. Two plant ESCRT proteins, AtVPS23A (ESCRT-I) and AtVPS4/SKD1 have been described to regulate plant cytokinesis^35^. How this proteins function in plant cytokinesis in detail is still unclear. It is plausible that the ESCRT function is essential in plants since loss of ESCRT results in lethality^46,51^. For example, the condensation of the ESCRT accessory protein FREE1 is necessary for a functional protein – otherwise, genetic knockout of *free1* in *Arabidopsis* results in seedling lethality^46,51^. Further experiments have to be done to analyse if HYA forms condensate-like structures. Previous reports showed that plant ESCRT components localized to endosomes and MVBs, and were required for the formation of ILVs within MVBs, and subsequently for the sorting of proteins for degradation^51–53^, In contrast, several lines of evidence indicate that HYA localizes to endosomal compartments and is involved in the secretory rather than the endosomal pathway (Fig. 6): first, *hya* mutants do not effect endocytosis; second, gHYA-GFP positive compartments are BFA and WM insensitive and show no colocalization with FM4-64 vesicles; third, in electron micrographs immunolabled gHYA-GFP was detected in PVCs; fourth, localization studies showed co-localization of gHYA-GFP with TGN derived compartments. Apparently, the presence of HYA in a plant-specific branch may indicate that the ESCRT-III component HYA, together with its ESCRT-interacting proteins, is involved in mechanisms other than those described for typical plant ESCRT components. We have demonstrated that gHYA-GFP accumulates during cell division around the newly formed cell plate and at the cortical division site. Moreover, gHYA-GFP was detected at the cell wall. In the past few years, several studies have brought to light substantial vesicle secretion during plant cytokinesis^5^. Whereas endocytosed plasma-membrane proteins could be targeted to the cell plate, only secretory trafficking of newly synthesized proteins is essential during cytokinesis^5^. Support for plasma membrane remodelling was provided by the observation that phospholipids like phosphatidyl-inositol-4,5-biphosphate [PtIns(4,5)P2] and phospholipid signaling moleclues accumulate at the leading edge of the cell^54–56^. The significantly reduced FM4-64 loading in *hya-3* indicates, either a different composition of the plasma membrane of the cell plate and/or less FM4-64 recycling events by exocytosis, we provide compelling evidence that HYA is part of some secretion mechanism during plant cell division that is necessary for plasma membrane remodelling. Further experimental evidence is required to determine the specificities of the ESCRT components in plants.

**Figure 6.**
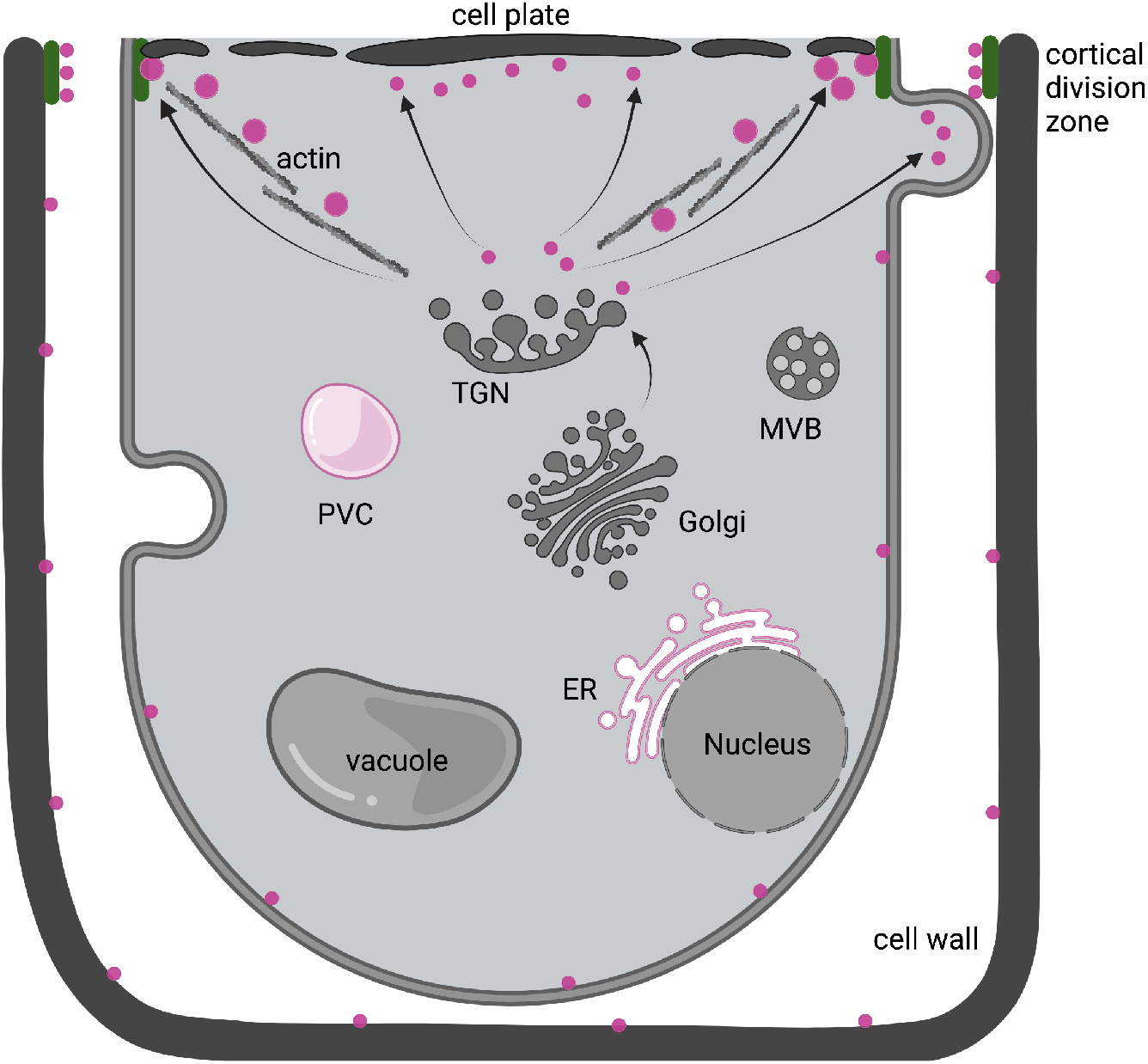
Schematic presentation of the localization and putative functional role of HYA. Schematic presentation of the localization and putative functional role of HYA (pink). TGN, transgolgi network; PVC, prevacuolar compartment; ER, endoplasmic reticulum; MVB, multivesicular body.

## METHODS

### Plant material, growth conditions and transformation

Plant material, growth conditions and transformation. Wild-type accessions Columbia (Col) and Wassilewskija (Ws) were obtained from the Arabidopsis Stock Center (Columbus, OH, USA). Provenance of the *hya-1*, *hya-2*, *hya-3* were described in^1^. *hya-4* was obtained from the NASC - European Arabidopsis Stock Centre (SALK059274;^57^. GFP-MAP4^22^ was a kind gift of H. Hö fte and D. Bouchez and pSNX1::SNX1-GFP from Y. Jaillais^39^. Plant growth conditions on sterile nutrient agar medium were described previously^58^. Plant transformation was done after the floral dip protocol of Clough and Bent^59^.

### Colocalization studies by transient transformation of mesophyll protoplast

For in vivo localization of HYA, a 2807 bp subfragment of pPZP211-gHYA was PCR amplified using primer 5g44560-GFP-SacI-pm and 5g44560-GFP-NcoI-R, and subcloned into pCR4-TOPO for sequence verification. Using SacI/NcoI the insert was transferred to the binary GFP-containing vector pGreenII0029-35S-GFP-RL replacing the 35S promoter. The final construct pmHYA::gHYA-GFP was transformed into *A. tumefaciens* strain GV3101(pMP90, pSoup) by electroporation and used to transform wild-type (WS, Col) and *hya-2*, *hya-3*. Transformants were selected on MS medium containing kanamycin (100 µg/ml). For the cloning of the 2×35S::mCherry-cHYA and 2×35S::YFP-cHYA fusions cDNA was amplified with primer HYA and cloned with SacI and BamHI into the pSAT6-mCherry-C1 and pSAT6-eYPF-C1 vectors, respectively. Similarily cDNAs of RabA3 (SacI and BamHI) and SYP61 (EcoR1 and BamH1) were amplified from leaf RNA and cloned into pSAT6-mCherry-N1 for the 35S::SYP61-mCherry construct and into pSAT6-mCherry-C1 for the 35S::mCherry-RabA3 construct. The RabF1/ARA6 construct 35S::ARA6-GFP was a kind gift of T. Ueda^47^. Rosette leaves of four-week-old soil grown plants were prepared, mesophyll protoplasts isolated and transformed as described by Sheen^60^.

### Phenotyping and microscopic analyses

GFP-MAP4 was crossed into *hya-1* and *hya-3* and segregating F2 and homozygous F3 seedlings were used for in vivo imaging of fluorescent markers in the confocal laser scanning microscope (CLSM, (TCS-SP2, Leica, Germany) via the Leica Confocal Software (Version 2.61) and a long working distance 63x water-immersion or a 100x oil objective. Excitation wavelengths were 488 nm (argon laser) for GFP and FM4-64 (SynaptoRed C2/FM4-64, Biotium, Germany). Emission for GFP was detected between 500 - 514 nm (GFP-MAP4) or 500-535 nm (gHYA-GFP) and between 635-680 nm for FM4-64. Pinhole and beam expander was set between one and three and the power to 25% (GFP-MAP4) and 48% (gHYA-GFP). For time series to follow cell division in vivo 1 picture min-1 was taken. The speed of the movies was accelerated 300-fold. All CLSM images were processed with the ImageJ program version 1.47n (http://rsb.info.nih.gov/ij/).

### FM4-64 staining and internalization

FM4-64 staining and internalization. To visualize the outlines of cells and quantify endocytosis, plasma membranes were labelled with a 8 µM aqueous FM4-64 (stock of 16.4 mM in aqua dest.) solution for 4 min at room temperature in the dark, rinsed with water and mounted after the indicated periods in water or low melting agarose media (LM; 1/2 Murashige and Skoog medium (MS), 1% saccharose) on slides with petroleum jelly border creating a chamber to accommodate whole seedlings.

### Inhibitor treatments

Inhibitor treatments. Four - to seven day old seedlings were treated in low melting agarose and half-strength MS-medium or water with 100 µM Brefeldin A (BFA) (stock of 1.8 mM in DMSO:EtOH, 1:1), 100 µM Wortmannin (WM) (10 mM stock in DMSO), 10 µM Oryzalin (20 mM stock in DMSO) or 10 µM Latrunculin B (20 mM stock in DMSO) and incubated at room temperature for the indicated times. pSNX1::SNX1-GFP seedlings were used as positive control for BFA and WM treatments. The movement of the FM4-64, gHYA-GFP and pSNX1::SNX1-GFP compartments was recorded every 20 sec and displayed in ImageJ 100-fold accelerated. Control treatments were performed with equal amounts of the respective solvents.

### Cloning of *HYA*

Genomic DNA was isolated from seedlings by a modified Cetyltrimethylammoniumbromid (CTAB) method (Hauser et al., 1998). Fine mapping with duplex analysis (mfc16not, mfc16sfi and 12D10R) and the restriction fragment length polymorphism (RFLP) (K15C-6.8) markers which corresponds to a 6.8 kb BamHI fragment of the bacterial artificial clone T6D12 (Hauser et. al 1998) narrowed the HYA region closer to the marker K15C-6.8 (Fig. 2a). Two overlapping PstI subfragments (6.8 kb and 15.7 kb) of BAC T6D12 containing the RFLP marker were subcloned into the binary vector pPZP211 and transformed via Agrobacterium tumefaciens (GV3101) in *hya-1* but failed to complement. Therefore, the adjacent gene At5g44560 was sequenced and revealed mutation in all hya mutants. The T-DNA insert in *hya-4* (SALK-059274) was identified on exon 6 of At5g44560 by PCR with primers mfc16sfi-F and LBa1 (Supplemental Table 1) and developed similar phenotypes. Finally, the identity of the HYA gene was confirmed by complementation of hya-2 with a genomic 3.3 kb fragment of the At5g44560 gene. This fragment consisting of 2141 bp coding-, 644 bp promoter-, 124 bp 5’, 175 bp 3’ untranslated- and 214 bp 3’ non-coding sequences, was amplified from genomic DNA with primers 5g44560-5Bam and 5g44560-3Bam and cloned into the BamHI site of the pPZP211 plasmid. The final construct (pPZP-gHYA) was verified by sequencing and transformed into *A. tumefaciens* (GV3101) by electroporation. Wild-type plants (Ws) and *hya-2* were transformed and selected on MS medium supplemented with 100 µg/ml kanamycin. Complementation was confirmed in the T2 generation by PCR with primers mfc16sfi-F and mfc16sfi-R that flank the deletion of the *hya-2* allele allowing the simultaneous detection of the transformed wild-type and *hya-2* allele in transgenic plants. The PCR fragments (230 bp wild-type, 200 bp *hya-2*) were separated on a 5% PAGE. Restoration of root growth and morphology in transgenic lines was observed under the stereomicroscope. For the HYA promoter fusion construct (pmHYA::GUS) a genomic fragment of 1220 bp containing sequences 645 bp upstream of the transcriptional start site and the coding region to the beginning of exon 2 was PCR amplified with the primers 5g44560-5Bam and 5g44560-GUS-BamHI-R, subcloned into the pCR4-TOPO for sequence confirmation and transferred via BamHI into the pPZP212-B3576 that contains the ß-glucuronidase gene, uidA.

### Electron microscopy and immunolabelling

Root tips were frozen in 150 mM sucrose using a Leica EMPACT (Leica Mikrosysteme GmbH, Vienna, Austria) high pressure freezer. Specimens were cryo-substituted at −80°C in acetone containing 1% OsO_4_ and 0.05% uranyl acetate in a Leica EMAFS according to^61^. Following a wash in ethanol, root tips were embedded through stepwise infiltration into LR-Gold (London Resin Company Ltd, Theale, UK) at 4° C and polymerized under UV light at 366 nm for 48 hours at −20°C. Ultrathin sections were processed for immunolabelling as follows: Blocking with 1% bovine serum albumin and 50 mM glycine dissolved in PBS (phosphate buffered saline) for 30 min; incubation in anti-GFP (Roche 11 814 460 001, Mannheim, Germany; 1:20 in BSA/glycine/PBS) for 1 h; four times wash in PBS for 10 min each; incubation in anti-mouse IgG labeled with 10 nm colloid gold (Sigma G7652, St. Louis, USA; 1:10 in BSA/glycine/PBS) for 30 min. For double immunolabelling, sections were incubated in a mixture of anti-GFP (1:10) and anti-VSRAt-1 (kind gift of Liwen Jiang; 1:10) for one hour followed by one hour treatment with a mixture of anti-mouse IgG labeled with 5 nm gold particle (Sigma G7427; 1:10) and anti-rabbit IgG labelled with 10 nm gold (Sigma G7402; 1:10). After rinsing in PBS and distilled water, samples were investigated in a LEO 912 TEM (LEO Electron Microscopy, Oberkochen, Germany) in zero-loss mode at an acceleration voltage of 80 kV.

### Modelling the three-dimensional (3D) structure of HYA and *hya-3*

Modelling the three-dimensional (3D) structure of HYA and *hya-3*. To determine the effect of the amino acid substitution on the 3D structure of the *hya-3*, HYA and *hya-3* were modelled to the 3D structure of CHMP3 (PDB-Code 2GD5) UCSF Chimera, production version 1.5.^32^. The 3D structures of the homo-hetero dimers were constructed with AlphFold^33^ and imaged via Chimera.

### Protein alignment and phylogenetic tree of the ESCRT-III complex components

The basic model of the ESCRT-III complex alignment and phylogenetic tree is based on published results^27^. For further analysis, following complete genomes, EST and protein datasets were searched for VPS2, VPS24, VPS20, SNF7, VPS46, and VPS60: *Ashbya gossypii* (Ag), *Cryptococcus neoformans var. neoformans* (Cn), *Chlamydomonas reinhardtii* (Cr), *Ostreococcus lucimarinus* (Ol), *Vitis vinifera*, (Vv) *Physcomitrella patens ssp patens* (Pp) at the NCBI database https://www.ncbi.nlm.nih.gov, *Lotus japonicus* (Lj) at the miyakogusa.jp database http://www.kazusa.or.jp/lotus/, *Dictyostelium discoideum* (Dd) at the Sanger Institute https://www.genedb.org/, and *Populus trichocarpa* (Pt) at the eukaryotic genomics database https://mycocosm.jgi.doe.gov/Phypa1_1/Phypa1_1.home.html. Additional database searches of VPS2 and VPS24 for *Danio rerio* (Dr), *Mus musculus* (Mm) and *Pan troglodytes* (Pt) were done at National Center for Biotechnology Information (NCBI). To visualize, alignments were done with Molecular Evolutionary Genetics Analysis (MEGA) version X^62^ using the ClustalW multi alignment. The evolutionary relationship was inferred using the neighbour-joining40 method and bootstrapping was performed with 1000 replicates. Sequence alignment results are visualized by Gendoc where black, dark grey and light grey shading indicate conserved amino acids of 100%, 80% and 60%, respectively.

### Histochemical analysis of GUS activity

GUS staining was done in 50 mM sodium citrate buffer, pH 7.0, 0.05% Triton X-100, 1 mM potassium ferricyanide, 1 mM potassium ferrocyanide and 1 mM X-Gluc (Roth, Germany) at 37 °C, 30 min (embryos seedlings) and 2 – 3 h (leaves, flowers). Staining was stopped and cleared with 75% ethanol. Seedlings were mounted on microscope slides in a modified Kaiser’s glycerol gel of 50% glycerol, 0.7% gelatine and 2.3% Lysoform (Henkel, Austria). Images were taken with a Nikon D70 digital camera mounted on an inverted Axiovert microscope (Zeiss, Germany).

### Yeast Two Hybrid Analyses

Total RNA of roots or cell suspension cultures were used for cDNA synthesis according to^63^. cDNAs were amplified with primers (Supplemental Table 1) suited for in frame cloning into the Y2H vectors pGADT7 and pGBKT7, inserted into pCR4-TOPO (INVITROGEN, USA) for sequencing and transferred to the two Y2H vectors. For interaction studies the yeast strain pJ69-4a was co-transformed with the AD and BD vectors by the PEG/LiCl heat shock method (Ito et al., 1983) and selected on synthetic drop out medium (SD, CLONTECH, USA) lacking Leu and Trp at 29°C. Protein-protein interactions were scored by spotting a liquid SD culture of the positive co-transformants on selective media lacking Trp, Leu and His (-L -T -H). The strength of interactions was determined by replating the co-transformants on SD/-T/-L/-H medium with increasing concentrations of the histidine analogue 3-amino-1,2,4-triazol (3-AT) (3, 8, 15 and 25 mM 3-AT, SIGMA-ALDRICH, Germany).

### Plasmolysis

After staining 7 day old seedlings with FM4-64 in LM media, seedlings were treated with and mounted in aqueous 0.5 M mannose solution for CLSM analysis.

## ACKNOWLEDGMENTS

We thank S. Neubert, Sonja Frosch and Romana Ranftl for technical assistance with the plant tissue cultures and the GUS-staining. We are obliged to U. Lutz-Meindl for her kind introduction and help at the electron microscope. We are grateful to C. Luschnig for sharing the pZP212-B3576 vector, L. Jiang for the anti-VSRAt-1 (VSR1) antibodies, D. Bouchez and H. Hö fte for the MAP4::GFP, K. Harter for the BiFC vectors, W. Rozhon and C. Jonak for the pGreenII0029-35S::GFP-RL vector, P.J. Hussey for the GFP::FABD line, T Ueda for the RabF1/ARA6-GFP construct, Y Jaillais for the pSNX1::SNX1-GFP, pSNX1::SNX1-mRFP lines, M. Menges, J.A.H. Murray and Bayer CropScience for the MM2d suspension culture, and G. Adam for the Y2H vectors and yeast strain. We also acknowledge the Nottingham Arabidopsis Stock Centre for distributing the SALK T-DNA insertion mutants, provided by J. Alonso and J. Ecker. We thank M. Grebe and C. Luschnig for discussion during the project. We thank J. Hilscher, C. Konlechner and A. Schweighofer for discussion and comments on the manuscript. This work was supported by grants from the FWF (P16410-B12, P17888-B14) to M.-T.H.

## AUTHOR CONTRIBUTIONS

Conceptualization, M.-T.H. and V.I.; methodology, M.-T.H., V.I., I.F., C.P. and S.M.; investigation, M.-T.H. and V.I.; writing-–original draft, M.-T.H. and V.I.; writing-–review & editing, M.-T.H., V.I. and S.M.; funding acquisition, M.-T.H; resources, M.-T.H. and S.M.; supervision, M.-T.H.

## Suppl. Figures

**Figure S1.**
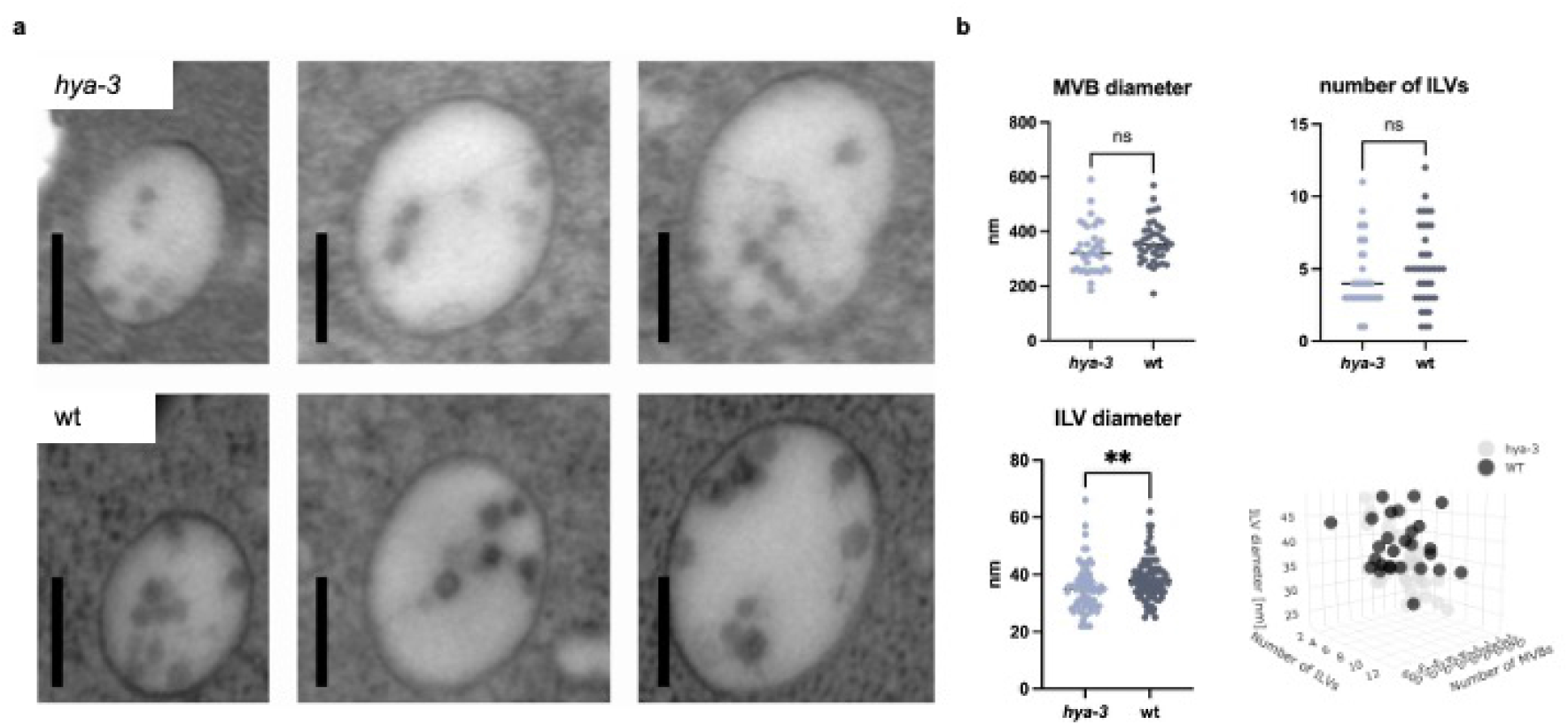
The morphology of MVBs in *hya-3* compared to wild-type. The morphology of MVBs in *hya-3* compared to wild-type. (a) MVBs in *hya-3* and in wild-type (wt). (b) The MVB diameter, the number of ILVs, and the ILV diameter was calculated. n = 29 wt, 38 *hya-3*. MVBs are cluster according to the average diameter of MVBs, ILVs and the number of MVBs. The scatter blot was blotted using R. Scale bars represent 0.2 µm.

**Figure S2.**
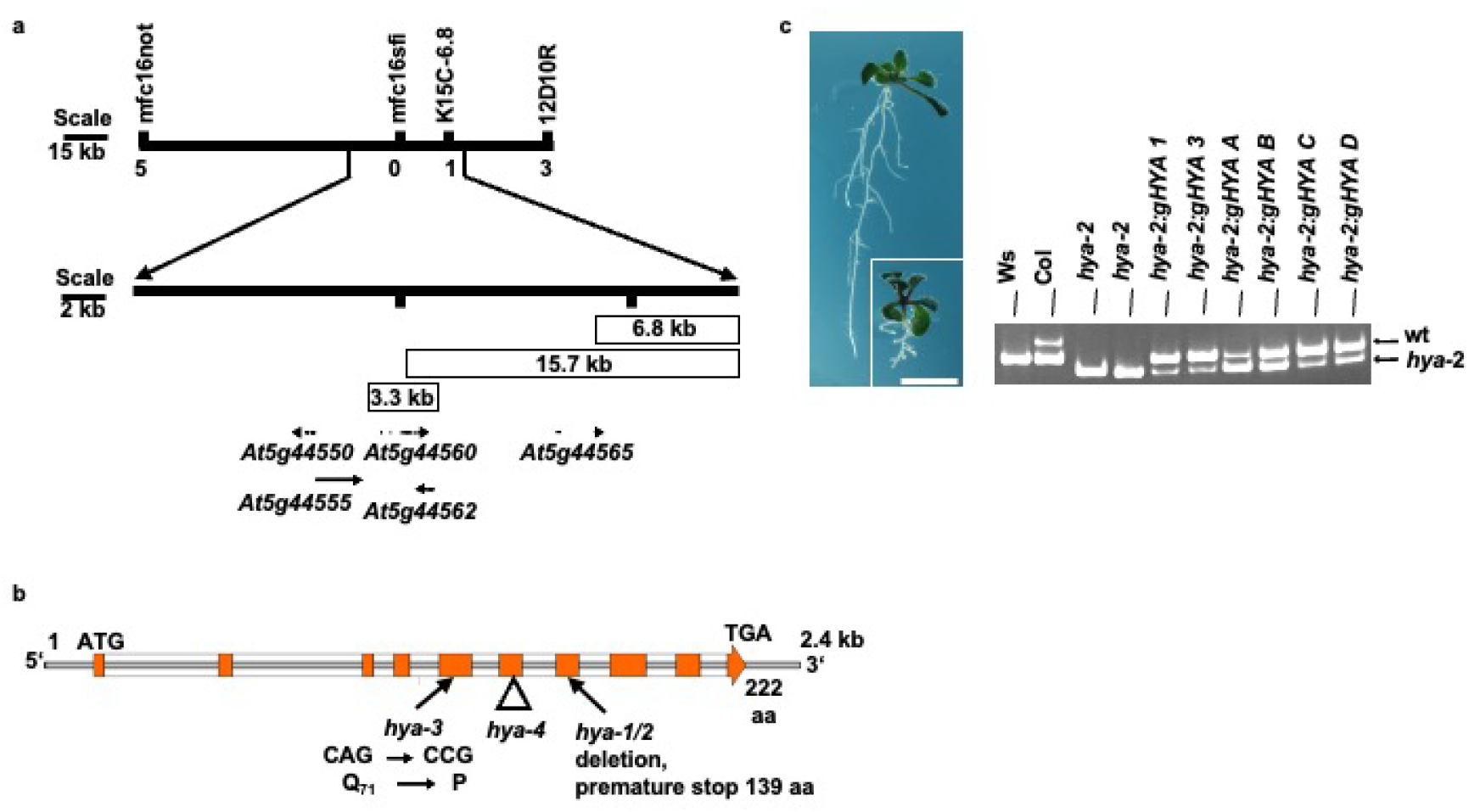
Isolation of *HYA/VPS2.2*. (a) *HYA* was mapped to chromosome 5 in a region of approx. 87 kb with a mapping resolution of 14.5 kb. (b) Mutations in *hya-1* to *hya-4*. (c) The presence of the wild-type and hya-2 alleles was confirmed by PCR with primers flanking the deletion of *hya-2* in the rescued transformants. While clones 6.8 kb and 15.7 kb did not complement, the 3.3 kb genomic region of gene At5g44560 rescued the *hya-2* phenotypes (a, c).

**Figure S3.**
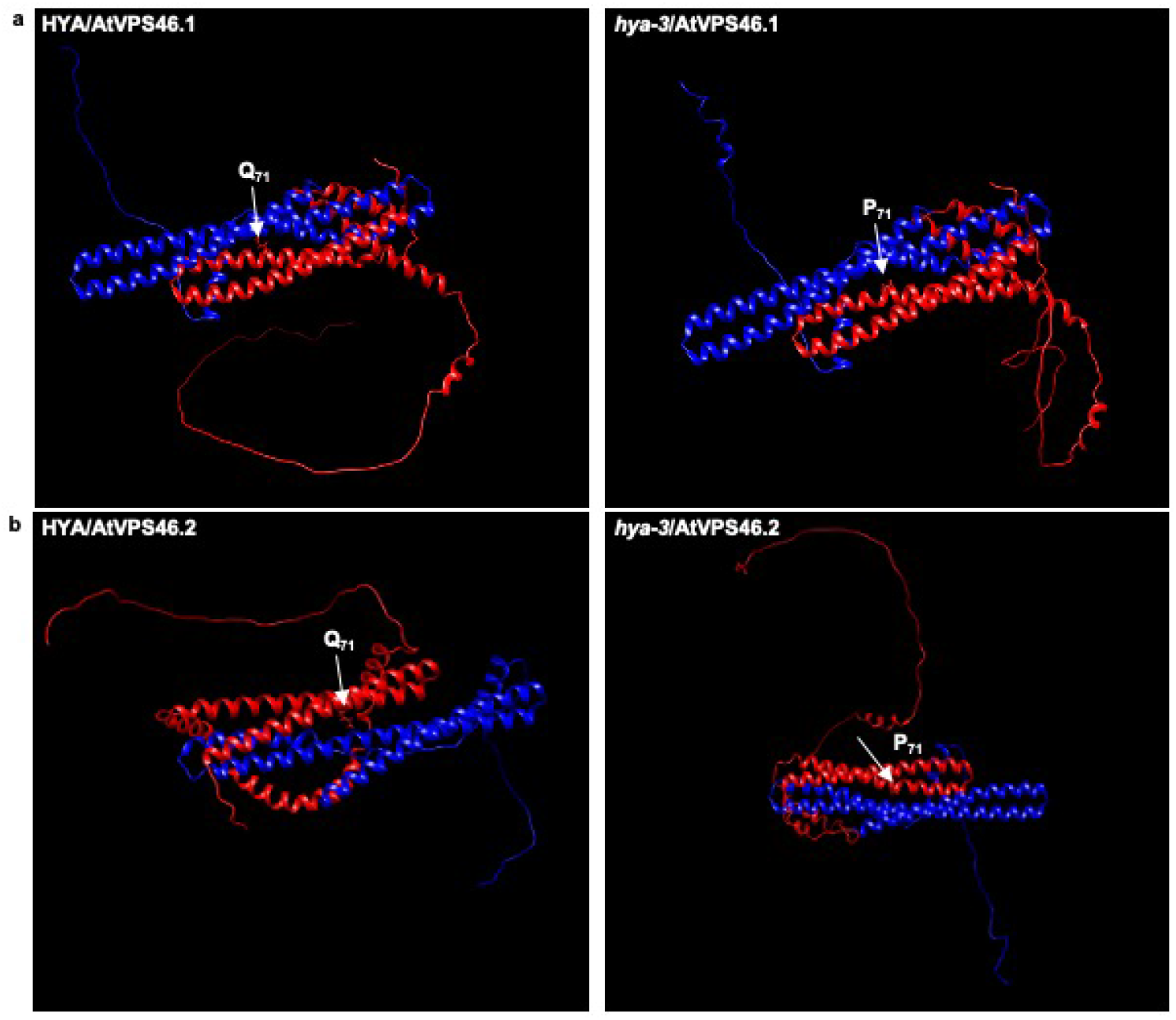
3-dimensional structural prediction of homo- and dimerization of HYA/AtVPS46.1 compared to *hya-3*/ATVPS46.1. 3-dimensional structural prediction of homo- and dimerization of (a) HYA/AtVPS46.1 compared to *hya-3*/ATVPS46.1 and (b) HYA/AtVPS46.2 compared to *hya-3*/ATVPS46.2.

**Figure S4.**
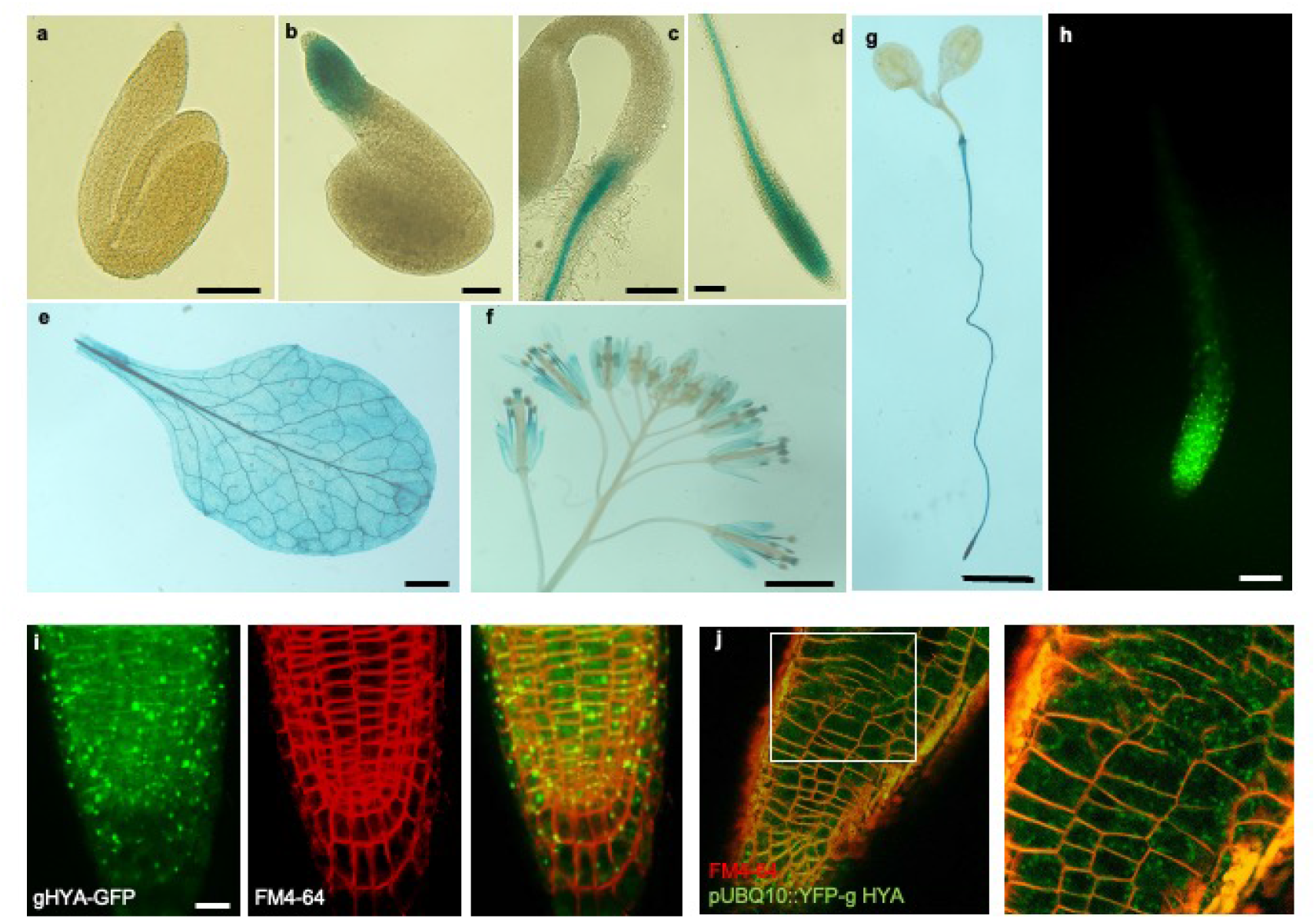
Characterization and complementation of HYA/VPS2.2 and *hya-3*. (a-h) Histochemical detection of pmHYA::GUS-and pmHYA::gHYA-GFP lines. HYA is developmentally regulated and not detectable in one day old seedlings of pmHYA::GUS lines (a). GUS activity was promptly apparent in root meristems two days after germination (b, c). In 4 days old seedlings the strong meristematic activity is maintained (c, d) and extents in the vasculature (c, d, g) of roots and mature rosette leaves (e). In the stigma, style and filaments of flowers GUS activity is only detectable after prolonged staining (f). (h) Meristematic activity of HYA was reproducible with the pmHYA::gHYA-GFP transgene. (i) *hya-3* was completely rescued by expression of pmHYA::gHYA-GFP. (k) *hya-3* was not rescued by expression of pUBQ10::YFP-gHYA HYA Scale bars equals 100 µm (a, b, d, h) 200 µm (c), 3 mm (e, f) and 5 mm (g).

**Figure S5.**
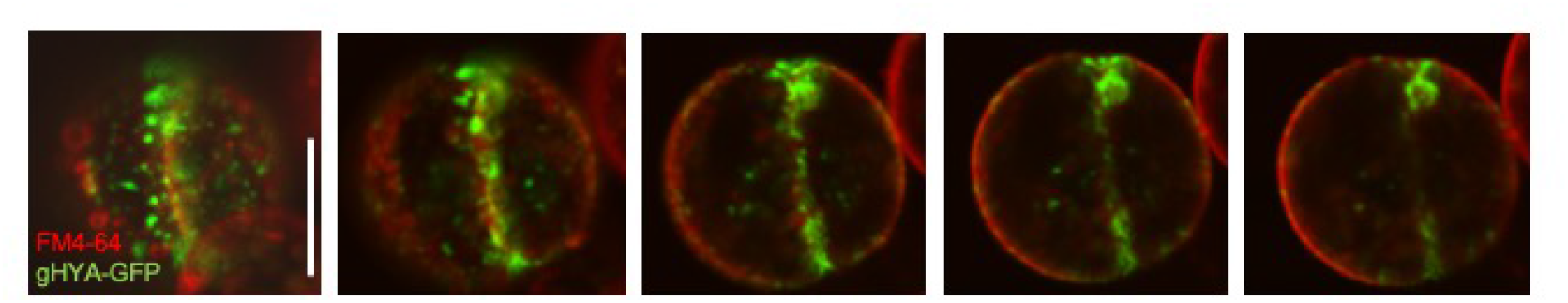
HYA localization during early cytokinesis in mesophyll protoplasts. gHYA-GFP transformed protoplasts were incubated over night at room temperature in darkness and stained in a 16 µM FM4-64 solution for 3 to 5 hours. Z-series are composed of 10 single scan images of 2 µm layer thickness. During late stages of cytokinesis HYA localizes around the cell plate and at the cortical division site. Scale bars equals 20 µm.

**Figure S6.**
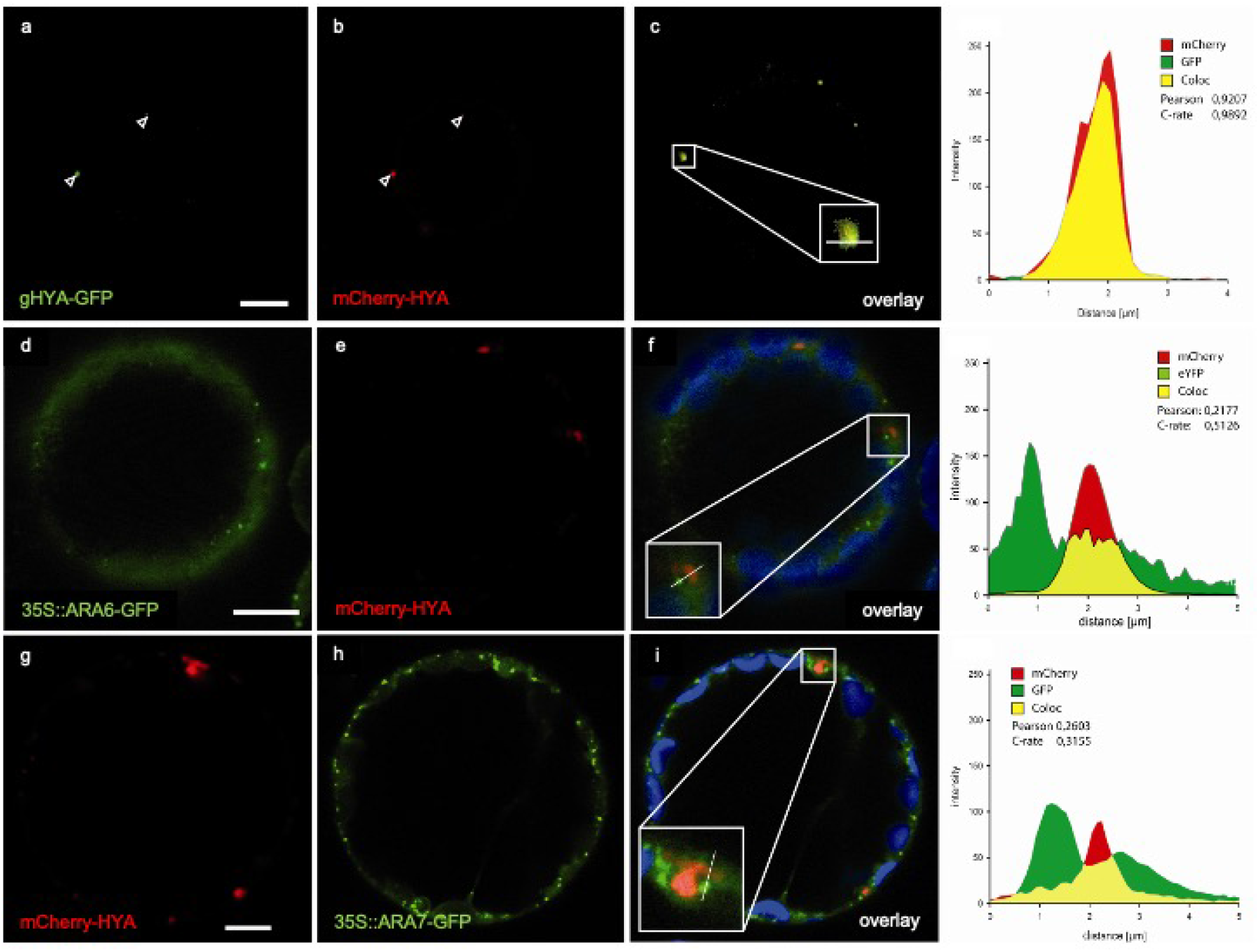
Interaction studies of HYA with endosomal markers in mesophyll protoplasts. Co-localization of gHYA-GFP with mCherry-HYA (a-c), no-colocalization of ARA6-GFP with mCherr HYA (d-f) and ARA7-GFP (g-i). Note the fluorescence profile of the overlays.

**Table S1.**
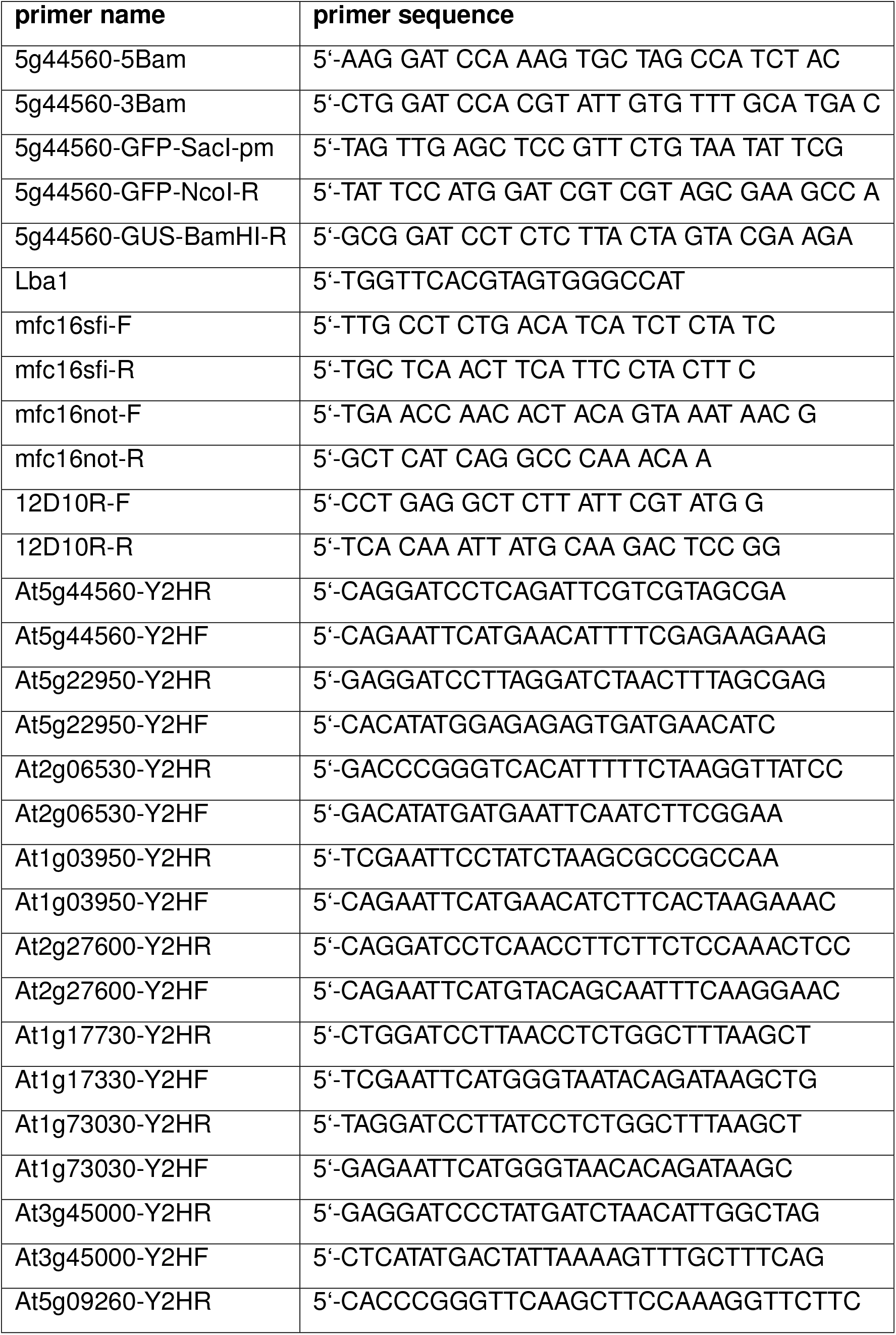
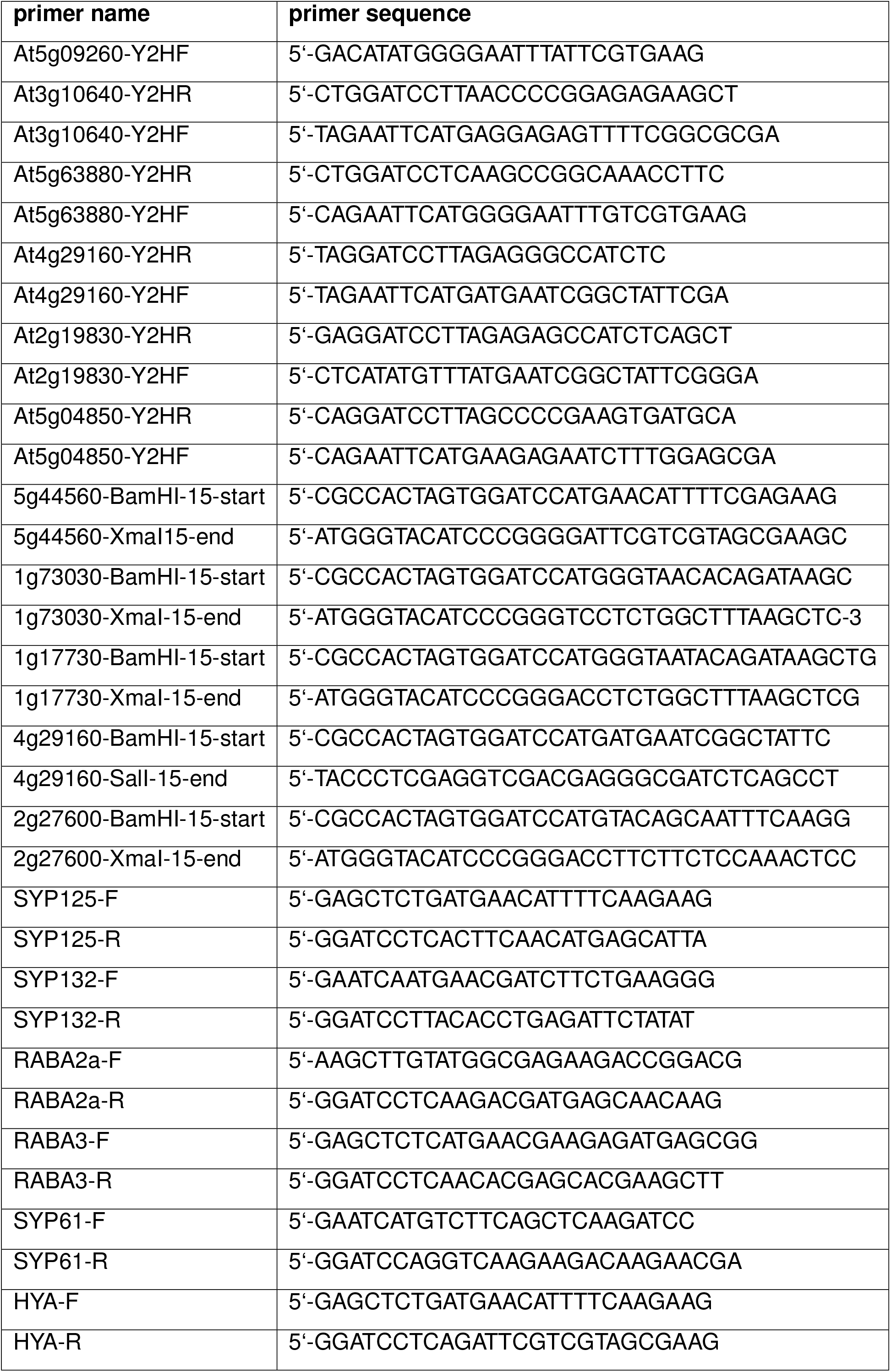
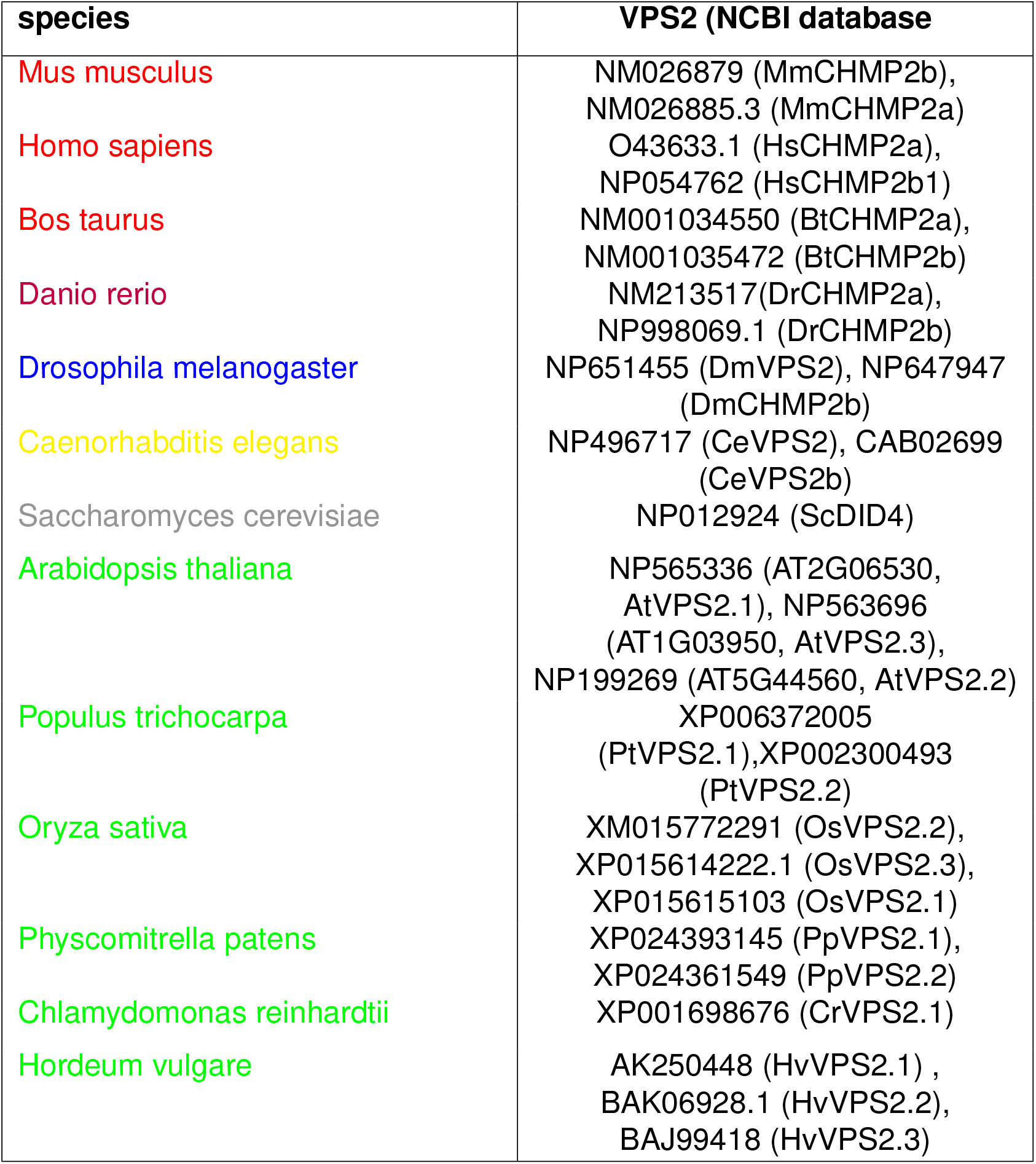
Overview of the primer pairs used for fine mapping and cloning of HYA plasmid construct for transcriptional and translational fusion, Y2H interactions analyses.

